# Functional Diversity of Memory CD8 T Cells is Spatiotemporally Imprinted

**DOI:** 10.1101/2024.03.20.585130

**Authors:** Miguel Reina-Campos, Alexander Monell, Amir Ferry, Vida Luna, Kitty P. Cheung, Giovanni Galletti, Nicole E. Scharping, Kennidy K. Takehara, Sara Quon, Brigid Boland, Yun Hsuan Lin, William H. Wong, Cynthia S. Indralingam, Gene W. Yeo, John T. Chang, Maximilian Heeg, Ananda W. Goldrath

## Abstract

Tissue-resident memory CD8 T cells (T_RM_) kill infected cells and recruit additional immune cells to limit pathogen invasion at barrier sites. Small intestinal (SI) T_RM_ cells consist of distinct subpopulations with higher expression of effector molecules or greater memory potential. We hypothesized that occupancy of diverse anatomical niches imprints these distinct T_RM_ transcriptional programs. We leveraged human samples and a murine model of acute systemic viral infection to profile the location and transcriptome of pathogen-specific T_RM_ cell differentiation at single-transcript resolution. We developed computational approaches to capture cellular locations along three anatomical axes of the small intestine and to visualize the spatiotemporal distribution of cell types and gene expression. T_RM_ populations were spatially segregated: with more effector- and memory-like T_RM_ preferentially localized at the villus tip or crypt, respectively. Modeling ligand-receptor activity revealed patterns of key cellular interactions and cytokine signaling pathways that initiate and maintain T_RM_ differentiation and functional diversity, including different TGFβ sources. Alterations in the cellular networks induced by loss of TGFβRII expression revealed a model consistent with TGFβ promoting progressive T_RM_ maturation towards the villus tip. Ultimately, we have developed a framework for the study of immune cell interactions with the spectrum of tissue cell types, revealing that T cell location and functional state are fundamentally intertwined.

---

Tissue-resident memory CD8 T (T_RM_) cells are a pivotal arm of the adaptive immune system, offering localized, long-term protection within non-lymphoid tissues through continuous tissue surveillance^1^. T^RM^ formation requires engaging transcriptional and metabolic programs to acquire tissue-specific adaptations^2–5^. These programs, initiated by priming events in lymphoid tissues^6^, are reinforced by cellular interactions and sensing environmental factors upon tissue entry^7,8^, such as TGFβ^9,10^, which enhances the upregulation of retention molecules including CD103 (*ITGAE*) in epithelial barriers such as the small intestine (SI)^9^. Newly infiltrated effector CD8 T cells responding to an infection in the SI display high interstitial motility, which becomes more restricted as cells become T_RM_ .^11^ Thus, signals received during the high-motility phase, such as short and long-range sensing of cytokines, might ultimately condition the destination of T_RM_ within the tissue. Recent studies have shown intestinal T_RM_ exhibit functional heterogeneity, with at least two distinct states identified in response to acute infections in the small intestine; an effector-like T_RM,_ and a polyfunctional memory T_RM_ population^3,12–14^ . These subpopulations display differential cytokine production and secondary memory potential, highlighting the nuanced nature of T_RM_ development and function that exist within tissues to provide long-term immunity. In fact, recent investigations into the architectural structure of the mouse and human small intestine have uncovered spatially organized transcriptional and metabolic programs that establish complex intestine regionalization^15,16^. How these tissue microenvironments affect T_RM_ differentiation and function has not been previously addressed. This is due in part to the fine granularity needed to capture different CD8 T cell transcriptional states and the variety of the non-immune cellular phenotypes that surround them, which requires highly multiplexed sub-cellular resolution imaging technologies. Spatial transcriptomics enables the profiling of hundreds of different messenger RNA molecules simultaneously within cells across complete tissue sections, in combination with protein and histology readouts. Here, we exploit spatial transcriptomics and develop a computational framework to characterize the cellular and ligand-receptor interactions guiding intestinal T_RM_ differentiation.

## A spatial framework for SI T_RM_

To quantitatively and systematically capture the spatial deployment of antigen-specific CD8 T cells in the small intestine responding to systemic LCMV infection, we used P14 T cell receptor transgenic CD8 T cells (P14 CD8 T cells), specific for the LCMV glycoprotein 33– 41 peptide (GP_33–41_) presented by H2-Db. After LCMV infection and adoptive transfer of P14 CD8 T cells from a CD45.1 congenic background into CD45.2 WT mice, small intestines (SI) were harvested at different times to analyze the number and intratissue location of transferred cells (**Extended Data Fig. 1a**). To obtain an unbiased assessment of intratissue location, we implemented a dual coordinate axes system based on the proximity of individual P14 CD8 T cells, detected by histological staining of CD45.1 and CD8a, to the nearest epithelial cell (x axis) or the distance to the base of the muscularis (y axis) (**Fig. 1a** and **Extended Data Fig. 1a**). This approach provided a 2D representation of the overall distribution of P14 CD8 T cells within the repetitive functional structure of the SI, which we termed IMmune Allocation Plot (IMAP) (**Fig. 1a**). IMAPs generated over the course of an LCMV infection captured the distribution dynamics of P14 CD8 T cells within the SI, revealing infiltration of the muscularis at an effector time point, followed by rapid clearance, and a subsequent formation of two spatially separated populations along the crypt-villus axis (**Fig. 1b**). These data suggest P14 CD8 T cells dynamically occupy different regions within the SI after infection.

**Figure 1.**
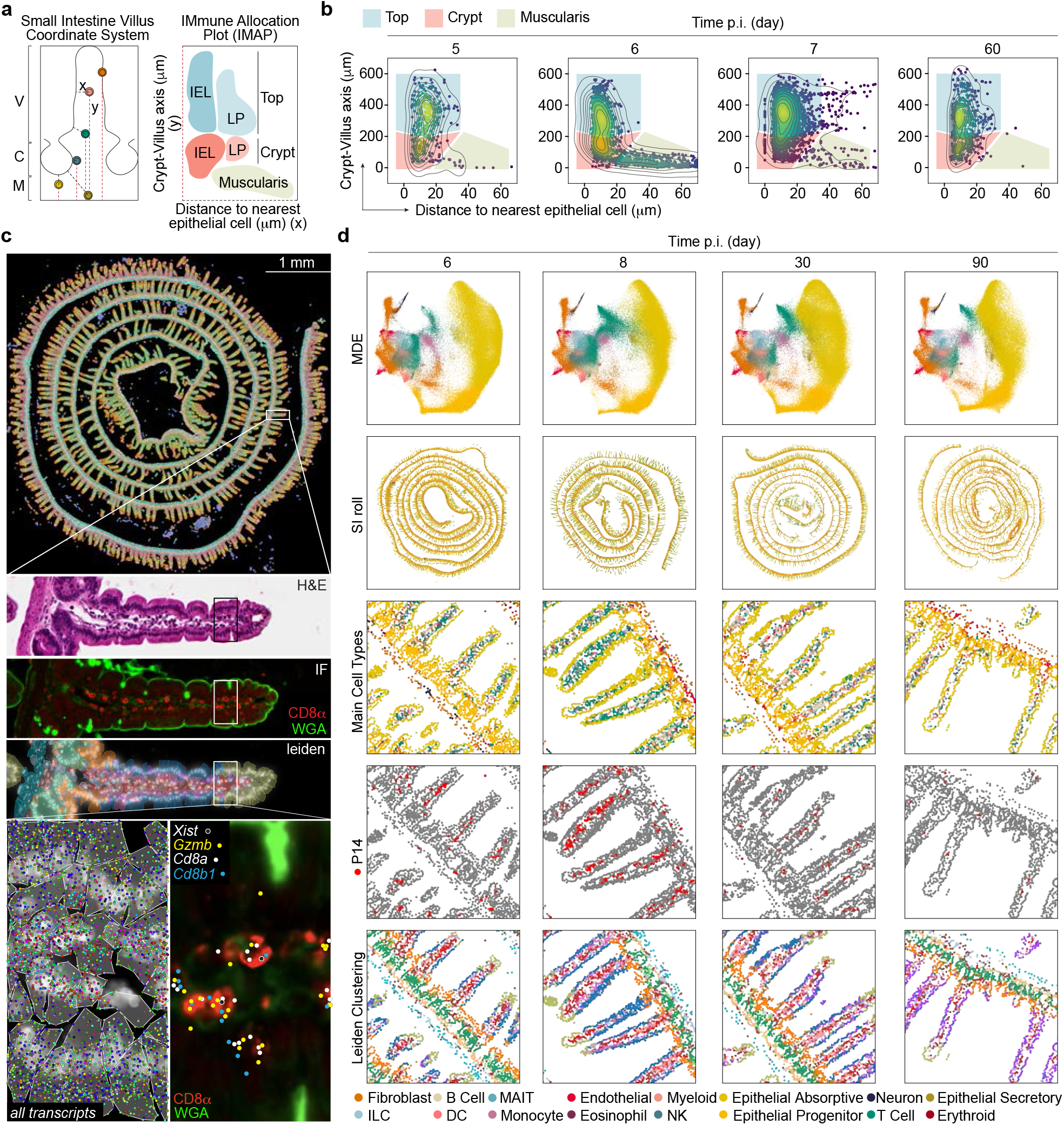
Characterization of the spatial and transcriptional state of antigen-specific CD8 T cells in responses to acute viral infection in the mouse small intestine with spatial transcriptomics. **a**, Diagram of a coordinate system to define morphological axes within the small intestine (left). Villus (V), Crypt (C), Muscularis (M) mark different physical regions in the tissue. Distance to the nearest epithelial cells (x) and distance to the base of the muscularis (y) form the basis of an Immune Allocation Plot (IMAP) (right). Top and crypt regions can house both intraepithelial lymphocytes (IEL) and lamina propria (LP) lymphocytes. **b**, IMAP representation of intestinal P14 localization progression measured by Immunofluorescence (IF) staining at timepoints post-infection (two biological replicates with n=3 mice per time point, 1 representative mouse shown). Gates for Top of Villus (Blue), Crypt (Red), and Muscularis (Beige) highlight the different regions. Points, representing cell positions, are colored by kernel density estimates over the IMAP (x,y) coordinates. **c**, Xenium-based spatial transcriptomics data structure overview. (row 1) Xenium output of a Day 8 pi mouse small intestine, with cells colored by Leiden cluster based on quantified RNA abundances. Zoom in of a villus showing (row 2) H&E staining, (row 3) confocal IF images of CD8α and plasma membrane marker Wheat Germ Agglutinin (WGA), and (row 4) Xenium DAPI staining with cell boundary segmentation masks overlaid and colored by Leiden cluster. (row 5) A further zoom in to a subregion of the same villus depicting (left) Xenium DAPI staining overlaid by cell boundary segmentation and all transcripts assigned to cells, and (right) IF image of CD8α and WGA with *Cd8a, Cd8b, Gzmb*, and female P14 specific *Xist* transcript locations overlaid. **d**, Processed Xenium data overview of mouse small intestines (SI) at day 6, 8, 30, and 90 takedown timepoints (columns). Rows depict (row 1) positioning of cells within a joint MDE embedding of all SI Xenium samples colored by cell type, (row 2) *in situ* spatial positioning of the cells, and close-ups colored by (row 3) cell type, (row 4) P14s highlighted red, and (row 5) Leiden cluster. One out of two biological replicates for each time point is shown.

To study the relationship of gene-expression differentiation programs of SI T_RM_ to anatomical location *in situ*, we performed spatial transcriptomic profiling (Xenium,10X Genomics)^17,18^ on SI from male mice adoptively transferred with female P14 CD8 T cells and analyzed over the course of LCMV infection (days 6, 8, 30, and 90). A 350-plex target gene panel was designed using a reference single nuclei RNAseq dataset to inform a prioritization algorithm based on predictive deep learning^19^ (**Extended Data Fig. 1b**). In addition, we included probes to detect relevant immune gene markers curated from the ImmGen database^20^, ligand-receptor pairs^21^, and *Xist*, a long non-coding RNA exclusively expressed in female cells that was used to track P14 CD8 T cells in the male hosts (**Extended Data Fig. 1b)**. Recursive feature elimination modeling showed 350 genes could capture the biological heterogeneity found in the SI (**Extended Data Fig. 1c, Supplemental Table 1**). After Xenium spatial transcriptomics, tissues were processed for immunohistochemical detection of CD8a and Wheat Germ Agglutinin (WGA), a pan cellular stain that enabled morphological segmentation of the villi structure, as well as H&E staining, which allowed assessment of overall tissue structures (**Fig. 1c**). Together, this analysis provided *in situ* measurements of 350 genes on 1.8 million cells over four time-points with biological duplicates (eight total SI samples), capturing an average median count of 98 transcripts per cell, and allowing identification of 36 distinct cell types, including bonafide CD8a^+^/*Cd8a*^+^/*Cd8b*^+^/*Xist*^+^ P14 CD8 T cells (**Fig. 1c and 1d and Extended Data Fig. 1e and 1f**). The gene-expression counts of the two biological replicates were tightly correlated (>95%) (**Extended Data Fig. 1d**). As expected with the course of infection, endogenous CD8 T cell and P14 CD8αβ T cell fractions were the most changed after LCMV infection with a notable increase in B cell frequencies (**Fig. 1d and Extended Data Fig. 1e**). Strikingly, spatial visualization of phenotypic diversity by Leiden clustering revealed a marked spatial pattern (**Fig. 1d**), suggesting our approach was able to capture transcriptional regionalization.

## SI T_RM_ diversity is spatially encoded

To study the transcriptional programs of SI T_RM_ as a function of their unique intratissue location, we generated IMAP representations for each cell type (**Extended Data 2a**) by calculating a coordinate along three main axes of reference for every cell: distance to the muscularis (crypt-villus axis) (**Fig. 2a**), distance to the nearest epithelial cell (epithelial axis) (**Fig. 2b**), and the position within the proximal-distal axis established along the gastrointestinal tract (longitudinal axis) (**Fig. 2c**). Crypt-villus versus epithelial axis IMAPs for P14 CD8 T cells across the infection timecourse revealed early accumulation in the muscularis, and progressive distribution up the intestinal villi, ending in two spatially segregated P14 CD8 T cell populations at day 90 (**Fig. 2d**), consistent with histology (**Fig. 1b**). P14 CD8 T cells were equally distributed along the longitudinal axis over time (**Extended Data Fig. 2b**). We next considered if gene-expression patterns were coordinated with spatial location for all cell types **(Fig. 2e and 2f)**. Out of the 191 genes detected in P14 CD8 T cells that were expressed in more than 5% of the cells, 87 (46%) had significant changes in expression along the crypt-villus axis, 76 (40%) changed along the epithelial axis, and 8 (4%) along the longitudinal axis (**Fig. 2e**). These results suggested strong transcriptional imprinting based on the intratissue location of P14 CD8 T cells, influenced primarily by the crypt-villus and epithelial axes. Other immune cells displayed similar correlations with variable expression along the epithelial and crypt-villus axis, while epithelial cells, most notably Enterocyte Subset 1, also showed a striking bias of correlated gene expression along the longitudinal axis for the interrogated genes (**Fig. 2f, Supplemental Table 2**), and in line with previous studies^15,22,23^. Genes associated with polyfunctional T_RM_ cells, *Tcf7* and *Slamf6*, and short-lived effectors (*Il18r1* and *Klrg1*), were expressed at the base of the villus in the crypt-villus axis, while genes associated with effector T_RM_ cells, such as *Itgae, Gzma* and *Gzmb* were highly expressed towards the top of the villus (**Fig. 2g and Extended Data Fig. 2c**). A similar transcriptional polarization was also observed along the Epithelial axis, with polyfunctional T_RM_ and short-lived effector cell associated genes preferentially expressed by P14 CD8 T cells in the LP (**Fig. 2g**). Unlike *Itgae*, which was stably expressed over time after infection, *Gzma* and *Gzmb* expression decreased concomitantly with an increase in *Tcf7* expression (**Fig. 2h and 2i**). Together, these data showed that CD8 T cell differentiation is transcriptionally controlled across time and space and suggested at least two different T_RM_ populations emerge late after infection. To put our results in the context of our previous observations, projection of polyfunctional T_RM,_ and effector T_RM_ gene signatures (clusters 3 and 29, respectively^13^) into the spatial coordinates of P14 CD8 T cells at day 90 of infection delineated a population of T_RM_ expressing cluster 3 genes preferentially located near the bottom of the villus, and T_RM_ enriched for expression of cluster 29 deployed at the top of the villus (**Fig. 2j**). Similar results were obtained using signatures derived from long-lived Id3^+^ T_RM_ and shorter-lived effector Blimp1^+^ T_RM_ cells^12^ (**Extended Data Fig. 2d**). Taken together, these data showed that T cells with a single TCR specificity (P14 CD8 T cells) show functional specialization associated with distinct cellular neighborhoods of the SI.

**Figure 2.**
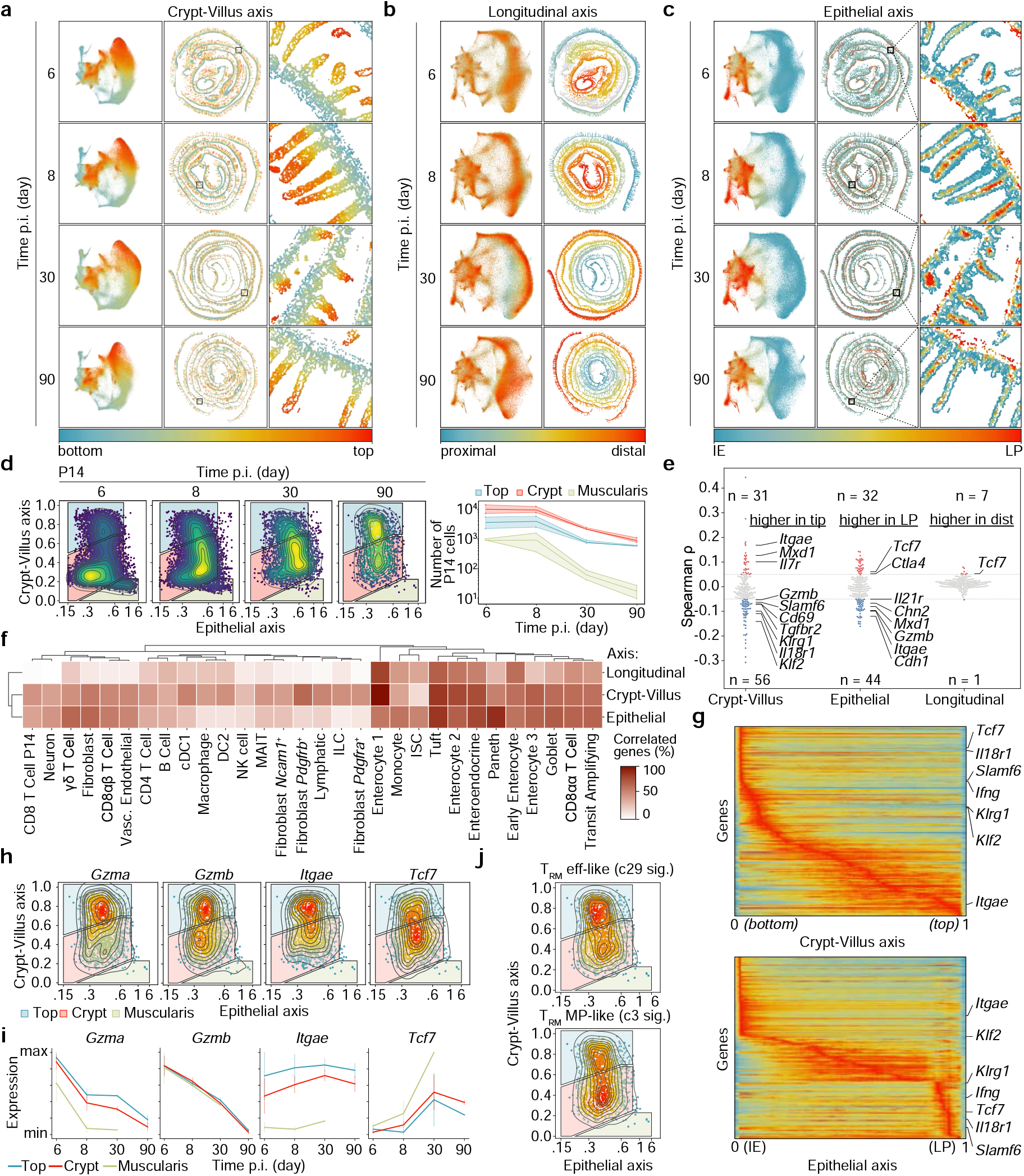
Intestinal regionalization along key axes instructs T_RM_ diversity in the mouse intestine. **a**-**c**, Spatial axes shown within the joint MDE embedding and *in situ* location, and **(a)** colored by their crypt-villus axis position from closer to crypt (blue) to closer to tip of villus (red), **(b)** colored by their longitudinal axis position from proximal (blue) to distal (red), **(c)** colored by their epithelial axis position from close to epithelial cells (blue) to closer to lamina propria (red). One out of two biological replicates for each time point is shown. **d**, (left) IMAPs of P14 CD8 T cells in samples from each time point, with colored gates dividing the top, crypt, and muscularis, and (right) the number of P14 CD8 T cells positioned within each regional gate across the time course (two biological replicates for each time point). **e**, Combined time course samples (n = 8), 4 time points with 2 biological replicates per time point, are pooled to create a swarm plot of Spearman rank correlation coefficients (ρ) between each axis and every gene expressed in at least 5% of P14s, with select correlated genes annotated. Genes are considered positively correlated (red) when ρ > 0.05, negatively correlated (blue) when ρ < -0.05, and not correlated (grey) otherwise. **f**, Heatmap depicting the percentage of genes in every cell subtype correlated with each axis using all combined time course samples (n=8). Heatmap colors indicate for all genes expressed in a particular cell type, few of them correlate with the corresponding axis (white), or most of them correlate with the corresponding axis (dark red). **g**, Convolved gene expression of P14 CD8 T cells along the crypt-villus axis (top) and epithelial axis (bottom) of all time course samples (n = 8). **h** and **i**, IMAP representations of day 90 P14s (one out of two biological replicates is shown) colored by kernel density estimates weighted by expression counts of select genes **(h)** and the expression of each gene within IMAP gated regions across time points in P14 CD8 T cells **(i)** (n=2). **j**, IMAP representations of day 90 P14 CD8 T cells (one out of two biological replicates is shown) colored by kernel density estimates weighted by UCell signature enrichment of polyfunctional memory T_RM_ (cluster 3^13^) and effector T_RM_ (“terminal state” (cluster 29)^13^). Signatures from Kurd et al. 2020. (LFC>1 and p-adj<0.01).

## Spatiotemporal progression of the T_RM_ interactome

To better understand how the anatomical position within the SI leads to T_RM_ heterogeneity over time, we focused on cellular interactions that were obtained by nearest neighbor analysis of our spatial transcriptomics dataset (**Fig. 3a**). Graphic representation of the cellular connectome of the SI separated four spatial domains: immune (LP), epithelial, muscularis, and crypt regions. Effector P14 CD8 T cells interacted mostly within immune cells of the LP and fibroblasts and over time, gained interactions with enterocytes (**Fig. 3b**). P14 CD8 T cells nearer to the tip of the villus (enriched for expression of cytotoxic molecules) were in closer proximity to enterocytes, and immune cells preferentially located in the top IE area, such as MAIT and NK cells (**Extended Data Fig. 2a**), while P14 CD8 T cells nearer to the crypts had increased physical proximity to B cells, CD4, fibroblasts and progenitor enterocytes (**Fig. 3c**). Thus, over time, P14 CD8 T cells gradually move away from the muscularis and diverge into a population that expresses higher levels of effector molecules that is in contact with differentiated enterocytes and immune cells of the villus top, while a T_RM_ population remains in the lower villus section where preferential interactions with CD4 cells, B cells, fibroblasts, and progenitor enterocytes are available. To capture the differential signaling along the villus, we focused on cytokine gradients along the crypt-villus axis captured in our Xenium dataset (**Fig. 3d**). Several key cytokines involved in T_RM_ formation and maintenance showed pronounced expression gradients, including *Il10*^24^, *Il7, Il21, Il15*^25,26^, and TGFβ isoforms^9^ (**Fig. 3d, e, f**). To obtain a more comprehensive view of cytokine gradients, and validate our findings, we profiled mouse small intestines at day 30 after LCMV infection using whole-genome spatial transcriptomics at 100 μm resolution (sequencing-based, Visium, 10X Genomics) (**Extended Data Fig. 3a**). While the resolution of this technology does not provide single-cell information, deconvolution of the crypt-villus axis using label-transfer information from our Xenium datasets revealed similar cytokine profiles (**Extended Data Fig. 3b and c)**. In addition, it uncovered cytokine expression gradients not previously included in our 350 gene panel, such as *Cxcl16* expressed at the top of the villus, which could mediate upper villus chemotaxis through its receptor CXCR6 (induced by TGFβ) on T cells. These data suggest that T_RM_ heterogeneity can be associated with differential access to key cytokines, including different sources of TGFβ ligands. Analysis of the differential incoming signaling patterns between P14 CD8 T cells located at the top or at the crypt highlighted MADCAM^27^, ICAM^28^, E-Cadherin, and TGFβ pathways as signals with differential patterns of expression associated with anatomical position, raising their potential role as upstream regulators of the heterogeneity observed in T_RM_ populations (**Fig. 3g and Extended Data 3d**).

**Figure 3.**
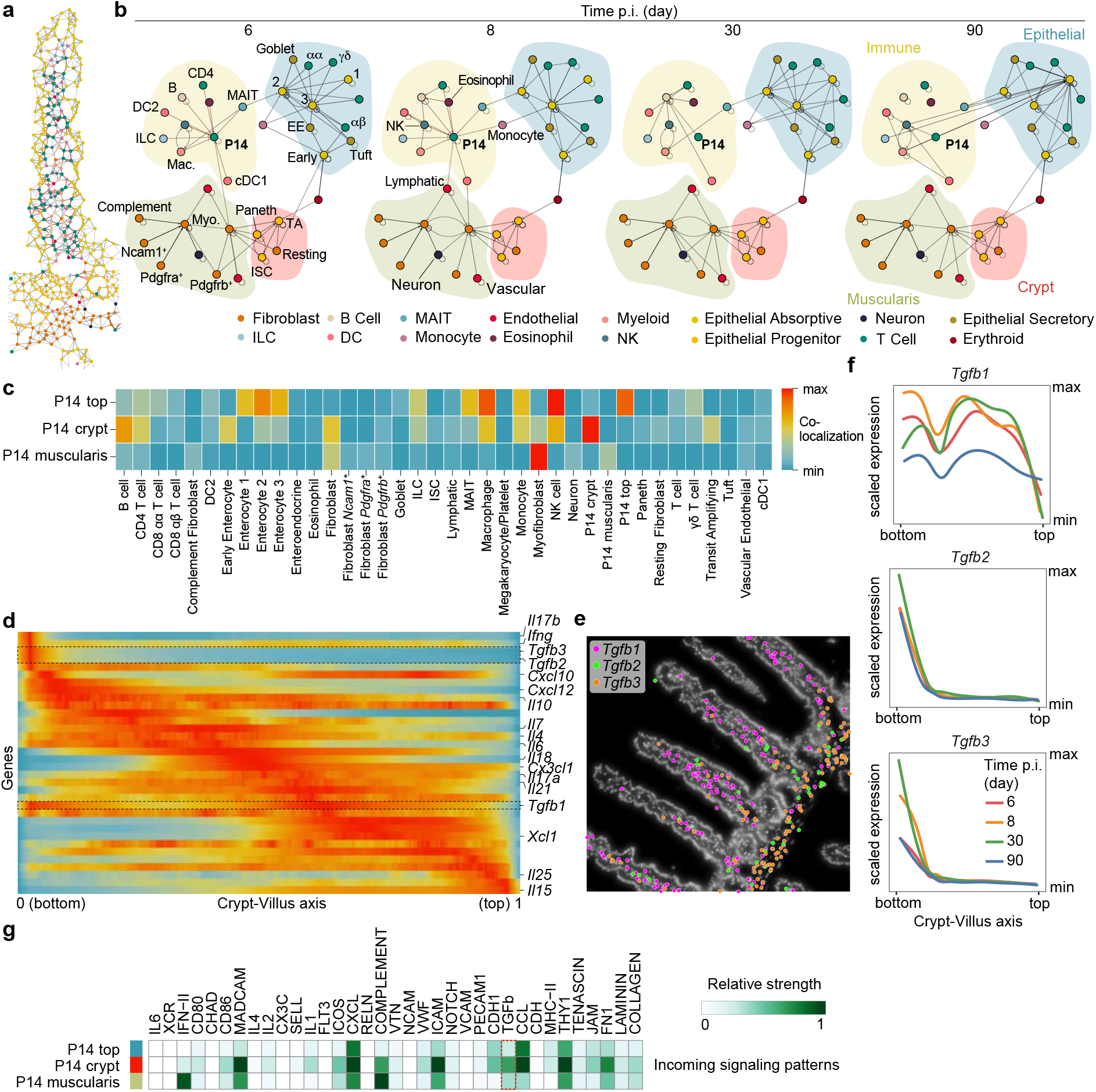
Cytokine gradients and differential interactions programs maintain T_RM_ diversity. **a** and **b**, Representation of the connectome between cell subtypes at (**a**) an individual cell resolution in a day 8 villus and (**b**) an aggregated network format where edges between nodes represent a normalized Squidpy interaction score lying above a 0.1 threshold (10% of the connections). Node (x, y) positions are determined by running a Kamada-Kawai layout algorithm on the Squidpy interaction matrix of the two day 8 replicates and visualized using igraph. For each time point, interaction scores between nodes are averaged across the two biological replicates. **c**, Squidpy interaction scores between cell subtypes and P14s regional groupings—Top, Crypt, and Muscularis as depicted in **Fig. 2d**. The color of the heatmap position reflects the strength of contact. Interaction scores are averaged across the eight samples, and values were row-normalized. **d** and **e**, Convolved gene expression of cytokines along the crypt-villus axis ordered and displayed with scVelo pooled across all time course samples (n=8). Isoforms of TGFβ are highlighted **(d)** and depicted spatially at their positions on villi from a day 8 pi SI **(e). f**, Gene expression trends for TGFβ isoforms separated by timepoint (n = 2 biological replicates pooled). A generalized additive model is used to fit a curve to the expression counts of each ligand along the crypt-villus axis, followed by standard scaling for comparison across trends. **g**, Heatmap showing the pathways contributing the most to incoming signaling of each P14 regional grouping. Relative strengths of each pathway were calculated using spatial CellChat on (n=8) samples from 4 time points. Heatmap is column-normalized across all subtypes, though only P14 CD8 T cells are displayed in the visualization.

## TGFβ sources contribute to programming of T_RM_ positioning

TGFβ is a powerful immunomodulatory cytokine that impacts intestinal CD8 T cell homing and retention^9,10^, and instructs T_RM_ differentiation and maintenance^3,29–31^. However, the sources of TGFβ isoforms, mediators, and cellular interactions that signal to CD8 T cells in the intestine are not well understood. Differential cellular communication analysis comparing P14 CD8 T cells at the villus base versus tip revealed a segregation of TGFβ isoform expression available to P14 CD8 T cells based on location (**Extended Data Fig. 4a**). Since P14 CD8 T cells showed similar expression levels of *Tgfbr1* and *Tgfbr2* within the villus (**Extended Data Fig. 4b**), we focused on exploring how cells of the SI might contribute to providing these signals. TGFβ is initially secreted as an inactive latent form that binds Latent TGFβ Binding Proteins (LTBPs) in the extracellular milieu. To signal through TGFβ receptors, the growth domain needs to be physically exposed via binding to αβ integrin dimers, such as alpha V (*Itgav*) coupled to beta 6 (*Itgb6*) or beta 8 (*Itgb8*), on the surface of trans-presenting cells^10^. Thus, the production and signaling of TGFβ can be decoupled. Consistent with this idea, expression of the TGFβ isoforms, as well as the presentation-associated molecules, suggested P14 CD8 T cells were exposed to TGFβ throughout their development, but the signaling might be provided by different cells based on anatomical location. For example, upper enterocytes (Enterocyte type 1) did not produce TGFβ isoforms but could, in theory, present TGFβ due to expression of *Itgav* and *Itgb6* (**Extended Data Fig. 4c**). To mechanistically explore this idea, we profiled the SI of mice that received WT or TGFβRII KO P14 CD8 T cells and were then infected with LCMV 8 days prior by spatial transcriptomics (**Fig. 4a**). To incorporate more elements of the TGFβ signaling pathway, we designed an expanded 494 gene panel and used the MERSCOPE platform (Vizgen), which utilizes MERFISH technology and includes three protein stains of cell boundaries (**Extended Data Fig. 4d**). This experiment generated a 470,000 cell dataset with an average of 186 transcripts per cell. TGFβRII KO P14 CD8 T cells, labeled by the expression of *Xist* and *Cd8* transcripts, had an overall spatial distribution shifted towards the bottom of the villi and muscularis area, and lower *Itgae* transcripts than WT P14 CD8 T cells, (**Fig. 4b and c**), while their total numbers were not changed at this time after infection, as previously observed^9^ (**Extended Data Fig. 4e)**. Differential gene expression between WT and TGFβRII KO P14 CD8 T cells showed global downregulation of the core T_RM_ signature^4^, as well as the TGFβ program (**Extended Data Fig. 4f**), with specific downregulation of *Itgae, Cxcr6, Cd160*, and *P2rx7*, and upregulation of *Klf2, Il18rap, S100a4*, and *Mki67*, consistent with published regulation by TGFβ signaling^32^ (**Fig. 4d**). Notably, *Mki67* was upregulated by TGFβRII KO P14 CD8 T cells and *Mki67*^+^ KO cells were differentially distributed in the tissue compared to WT cells, suggesting loss of adequate TGFβ programming caused misslocalized hyperproliferation closer to the crypts (**Fig. 4e**). To understand how P14 CD8 T cells engage in the TGFβ program based on their proximity to other cell types, we used the 8 differentially expressed genes as a TGFβ signature, which was correlated to the relative distance of each P14 CD8 T cell to every other cell type (**Fig. 4f and g**). The distribution of WT P14 CD8 T cell values showed that WT P14 CD8 T cells had the highest expression of the TGFβ program when closest to Enterocytes, and other cell types positioned in the top part of the villus, and lowest when WT P14 CD8 T cells were in the proximity of cells located in the muscularis (fibroblasts), crypt, or lower villus area (**Fig. 4g**). TGFβRII KO P14 CD8 T cells had a global loss of transcriptional-distance correlation, consistent with the notion these cells no longer sense TGFβ signals (**Fig. 4g**). Gene-expression visualization of TGFβ isoforms and components of the trans-presentation machinery showed fibroblast populations to be producers, consistent with our Xenium dataset (**Fig. 4g, Extended Data Fig. 4c**). Next, we analyzed differences in the physical distances between WT and TGFβRII KO P14 CD8 T cells to each nearest cell type. These analyses showed an increased accumulation of TGFβRII KO P14 CD8 T cells around fibroblasts while they were farther away from enterocytes and other cells located at the top of the villus than WT cells (**Fig. 4g and 4h**). These changes were not explained by the compensatory expression of TGFβ molecules (**Extended Data Fig. 4g**). Thus, P14 CD8 T cells obtain initial TGFβ programming by fibroblasts located lower in the villus, programming their positioning at the top of the villus as they engage the TGFβ program, where it can be further maintained by TGFβ-presenting cells, such as enterocytes (**Fig. 4i**).

**Figure 4.**
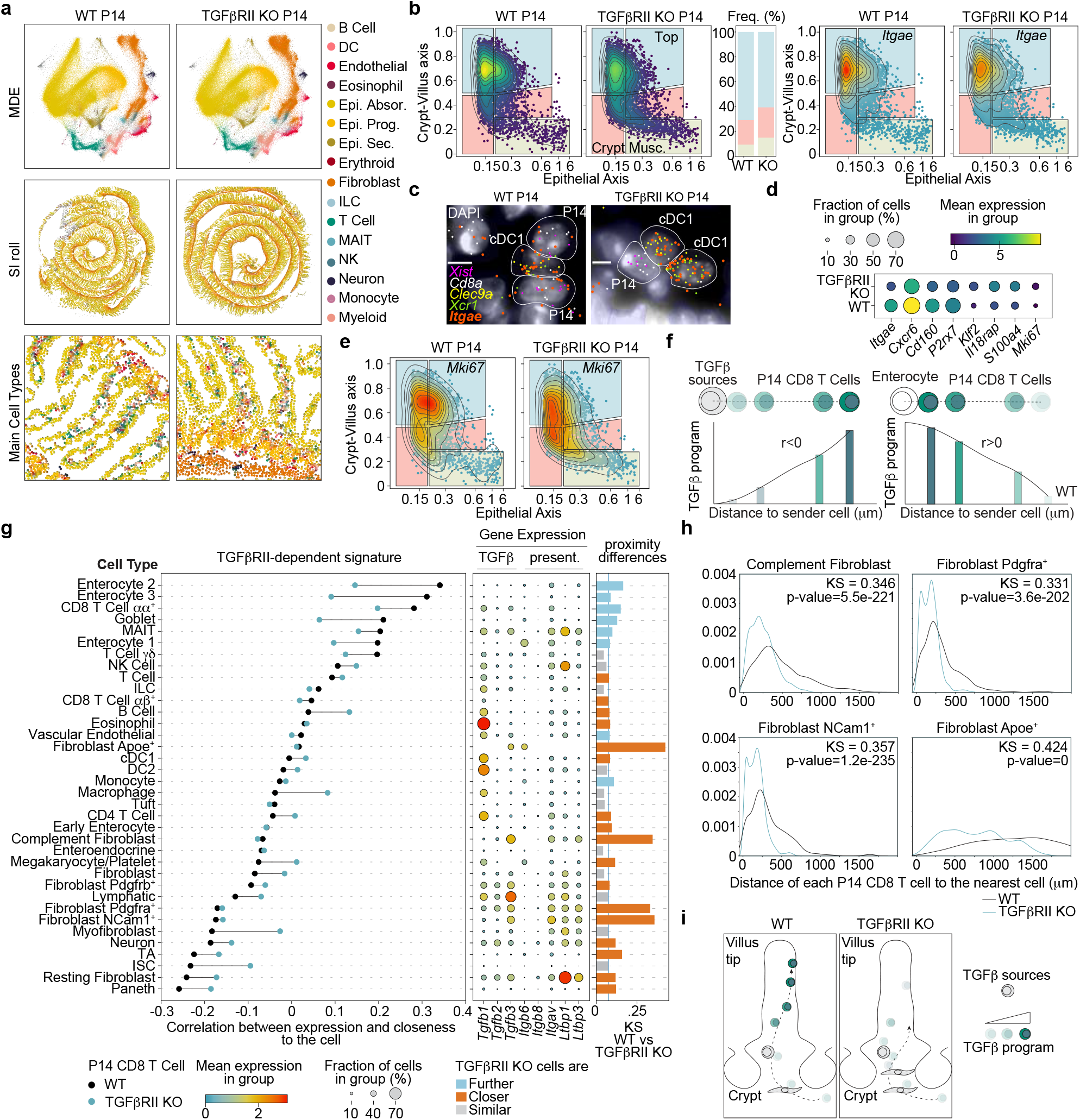
Sources and mediators of the TGFβ program for CD8 T cells in the SI profiled by MERSCOPE. **a**, Female wild type or TGFβR2 KO P14 cells were transferred into male C57BL/6 recipients. MERSCOPE-based spatial transcriptomics of the SI was done on day 8 post-infection. One WT and one TGFβR2 KO SI were profiled from one biological replicate with 3 mice per condition. Plots show the joint MDE embedding colored by cell type (top), *in situ* spatial positioning of the cells (middle), and close-ups (bottom). **b**, (left) IMAP positioning and kernel density estimate (weighted by gene expression, bottom) coloring of WT and TGFβR2 KO P14 cells. Cells are gated into top intraepithelial, top lamina propria, crypt intraepithelial, crypt lamina propria, and muscularis.(middle) Quantification of P14 cells localized in the muscularis, crypt or top of villus. (right) IMAP colored by kernel density estimates weighted by expression counts of *Itgae*. **c**, Contacting P14s and conventional dendritic cells in zoomed-in regions of the SI. DAPI staining is overlaid with scattered points representing the positions of select transcripts. **d**, Top differentially expressed genes between WT and TGFβR2 KO P14 cells. The dot plot is colored by the mean expression of each gene, and the dot size reflects the percentage of P14 cells in which the corresponding gene is expressed. **e**, IMAP representations of WT and KO P14 cells colored by kernel density estimates weighted by expression counts of the proliferation marker *Mki67*. **f** and **g**, TGFβR2 dependent signature enrichment and distance to each cell subtype were calculated for all P14 cells, and Spearman rank correlated against each other **(f)**. (**g**) For every subtype, (left) the correlation coefficients between signature enrichment and P14 cell proximity to the subtype among both WT and TGFβR2 KO P14 CD8 T cells, (middle) the expression of TGFβ isoforms and genes involved in TGFβ presentation in the WT sample, and (right) a non-parametric Kolmogorov– Smirnov statistic indicating the significance of difference of the distance distributions between P14 CD8 T cells and the corresponding cell type in both WT and TGFβR2 KO. The color of the bars indicates whether P14 CD8 T cells are closer to a given cell type in WT (blue) or TGFβR2 KO (red), and a line indicating effect relevance is positioned at 0.08. **h**, Comparisons of the distance between WT or TGFβR2 KO P14 cells and selected other cell subtypes. A Kolmogorov–Smirnov statistic indicates the difference between the WT and KO distributions for each subtype. The plotted lines show the positional density using a 1D kernel density estimate. **i**, Proposed mechanism of effector T_RM_ maturation in the SI, where fibroblasts use TGFβ signaling to prime antigen-specific T-cells into the molecular programs needed for residence at the villus tip.

## Spatial imprinting of human SI CD8 T cell diversity

To put the relevance of these findings into the context of human intestinal immunity, we profiled two terminal ileums from healthy donors in technical replicates using the Xenium platform with a custom immune-focused 422 gene panel (**Extended Data Fig. 5a** and **Supplementary Information Table 3**). Combined, we generated a dataset comprising 214,546 cells with an average of 89 median transcripts per cell and identified 38 different cell types (**Fig. 5a and Extended Data Fig. 5b**). Technical replicates (consecutive slides) were nearly identical, and the observed cell frequencies were similar between biological samples, except for B cells due to differences in the abundance of Peyer’s Patches (**Fig. 5b and Extended Data Fig. 5c**). IMAPs were computed by calculating the crypt-villus axis using spatial transcriptional neighborhoods around each cell as a predictor for distance to the basal membrane, and epithelial distance axes using the normalized distance to the nearest epithelial cells (**Fig. 5c**). First, to explore whether spatial distribution differences were observed across T cell populations, we focused on T cell types that emerged from unsupervised clustering (**Extended Data Fig. 5d**). CD8 T cells, proliferating T cells, and γδ T cells were predominant in the top IE area, while effector T cells, including *GZMK*^+^ CD8 T cells, were predominantly localized in the LP and more evenly distributed along the crypt-villus axis (**Extended Data Fig. 5e**). These differences in location were reflected by their distinct interactions with nearby cells (**Extended Data Fig. 5f**). Next, we focused our analysis on *CD8A*^+^*CD8B*^+^*CD3G*^+^*CD3D*^+^*CD4*^-^ T cells (CD8 T cells) outside of the Peyer’s patches (**Fig. 5c**). Projection of the transcriptional signatures from the two spatially segregated P14 CD8 T cell populations we observe at day 90 in mice identified similarly segregated subpopulations in human CD8αβ T cells (**Fig. 5d**). Furthermore, spatial differential expression analysis between human CD8αβ T cells in different morphological regions recapitulated mouse findings, as many genes are significantly spatially regulated, often similarly to their mouse homologs. These include GZMA and ITGAE which localize near the epithelium at the top of the villus, and KLRG1 and TCF7 which localize to the LP and bottom of the villus as they do in mouse P14 CD8 T Cells. (**Fig. 5e and 5f, and Extended Data Fig 5g)**. These data show that the spatial imprinting of the phenotypic and functional diversity observed by mouse CD8 T_RM_ cells occurs similarly in humans. Analysis of gene expression dynamics as a function of the relative distances of CD8 T cells to other cell types highlighted effector molecules ITGAE and GZMA to be most highly expressed by CD8αβ T cells when in proximity to enterocytes and long-lived memory associated molecule TCF7 to be most highly expressed by CD8αβ T cells when in proximity to CD4 and B cells (**Fig. 5g**). Finally, differential cellular communication analysis based on spatial distances and ligand-receptor expression of polyfunctional Id3, TCF expressing T_RM_ cells and effector T_RM_ (defined by the mouse spatial signatures, as in **Fig 5d**) identified differential incoming signaling (**Fig. 5h**). Among these, ICAM (communication with Vascular Endothelial) was more prevalent in the memory-signature enriched CD8αβ T cells, while C-type lectins (CLEC), CC chemokine signaling (CCL, communication with other immune cells) were favored for effector-signature enriched CD8αβ T cells and TGFβ signaling was similar for both CD8αβ T cell subtypes (**Fig. 5h**). In sum, these data suggest that the heterogeneity of phenotype and gene expression observed by CD8 T cells in the small intestine is imprinted by their strategic intratissue location, especially along the crypt-villus axis, which, through differential cellular interactions and exposure to cytokine sources, such as TGFβ, maintains functionally different populations of CD8αβ T cells in both human and mouse.

**Figure 5.**
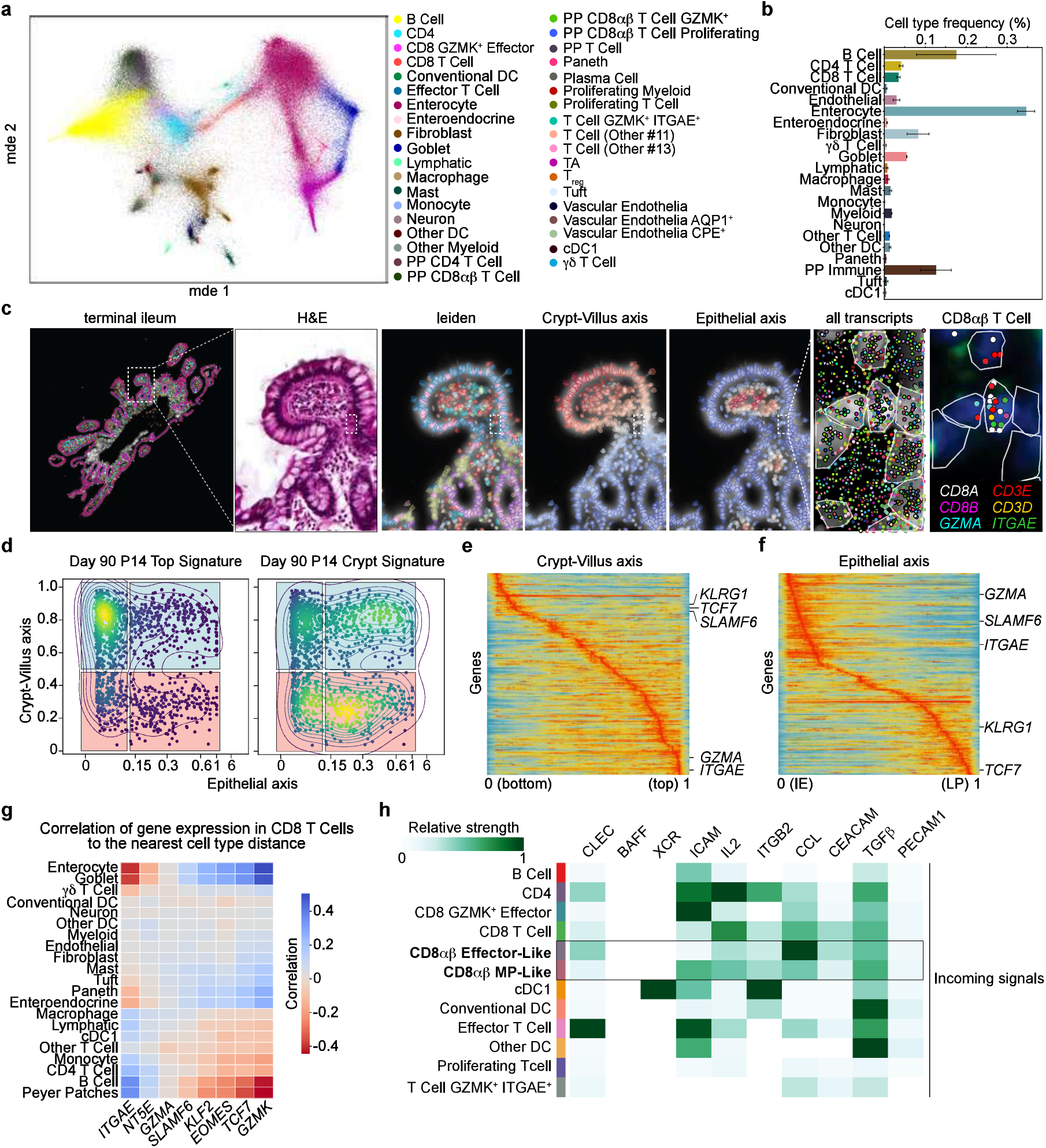
CD8 T cell phenotypic diversity in the human ileum is spatially imprinted. **a** and **b**, Spatial transcriptomics of human small intestine sections from two donors (2 consecutive tissue sections from each) using 10X Xenium. (**a**) Joint MDE embedding by cell type and (**b**) the mean relative frequencies of each annotated cell type across all sections pooled. Error bars represent the standard error of the mean across (n = 4) human sections. **c**, Overview of the Xenium-based spatial transcriptomics data from the submucosal biopsy of human terminal ileum. From left to right, (1) Xenium output of a human terminal ileum, with cell segmentation masks colored by annotated cell type. Zoom in of a villus showing (2) H&E staining, Xenium DAPI staining with cell boundary segmentation masks overlaid and colored by (3) Leiden cluster, (4) Crypt-Villus Axis, and (5) Epithelial distance. A further zoom-in to a subregion of the same villus depicting (6) Xenium DAPI staining overlaid by cell segmentation masks and all detected transcripts and (7) select transcripts overlaid upon DAPI staining. **d**, IMAP representation of human CD8αβ T cells colored by kernel density estimates weighted by the mouse signature for P14 cells at the top of the villus (left) or in the crypts (right). Human IMAP gates define the top of the villus (blue) and crypt (red) and split down the middle to define intraepithelial (left) and lamina propria (right). Human CD8αβ T cells are pooled across all human samples (n=2 donors, 2 adjacent tissue sections from each), and Peyer’s Patches are excluded. **e** and **f**, Pooling all human samples (n=2 donors, 2 adjacent tissue sections from each), and excluding Peyer’s Patches, convolved gene expression of CD8αβ T cells along the (**e**) crypt-villus axis and (**f**) epithelial axis. **g**, Expression of select genes in CD8 T cells are Spearman rank correlated with distances between CD8 T cells and other cell types. A red color indicates that P14-specific expression of a gene is increased when T cells are near a given cell type (expression is negatively correlated with increasing distance). Conversely, a blue color indicates that the expression of a gene is decreased when T cells are near the corresponding cell type. Correlations were calculated for each sample (n=4) individually and the mean correlation coefficient is shown. **h**, Heatmap showing the most contributing pathways to incoming signaling of different human immune cell groupings. Relative strengths of each pathway were calculated using spatial CellChat on all human samples. Heatmap is column-normalized across all human cell subtypes, though only specific immune subtypes are displayed in the visualization. Human CD8αβ T cells were grouped either effector or stem-like based on their enrichment of the mouse-derived top and crypt UCell signatures. Enrichment of each signature was z-scored across all human CD8αβ T cells before direct comparison to classify the CD8αβ T cells into effector-like or stem-like groupings.

## Discussion

High-resolution spatial transcriptomic interrogation of mouse models of viral infection and human samples revealed that the internal intestinal cellular architecture shapes CD8 T cell diversity in the intestine. Our findings support a model in which the cellular organization of the intestine provides localized instructions through regionalization of cytokine secretion and distinct cellular interactions that lead T_RM_ differentiation from the lower villus to the upper intraepithelial area, where T_RM_ are exposed to cytokines that promote their persistence. In parallel, spatial patterning throughout the crypt-villus and epithelial distance axes provide at least two different T_RM_ interactomes that lead to the formation and maintenance of two functionally and phenotypically distinct populations of T_RM_, observed both in mouse and human, which include T_RM_ in the tip of the villi that express cytotoxic effector molecules and a long-lived, polyfunctional T_RM_ subset in the lamina propria at the base of the villi.

Our data provide a new understanding of how the TGFβ program is imparted on intestinal CD8 T cells to promote T_RM_ programs *in situ*. While many cells, both in mice and humans, can express TGFβ isoforms (**Extended Data Fig. 5h)**, loss of TGFβRII leads to a preferential accumulation of CD8 T cells around fibroblasts in the lower villus area and separation from upper enterocytes. This suggests that fibroblasts provide an initial TGFβ signal needed to direct pre-T_RM_ cells to areas where they will be exposed to subsequent signals that promote further maturation. Of note, while enterocytes are not producers of TGFβ in mice or humans, they express the integrin (Integrin Alpha V, *ITGAV*) required for the presentation of the active TGFβ signal (**Extended Data Fig. 5h**). In the skin, migratory DCs use Itga_v_ to provide TGFβ stimulation to naive CD8 T cells to precondition them into skin T_RM_ fate, while keratinocytes maintain epidermal residence^10,30^. In the future, it will be interesting to determine to what extent TGFβ presentation by specific cell types is needed to maintain IE CD8 T cell populations.

Our *in vivo* approaches and computational analysis to systematically study the spatial positioning of T_RM_ within the small intestine could also be applied to other tissues with functional repetitive structures, such as the nephron in the kidney, glandular structures, or the hepatic lobule and similarly provide a framework for the study of other immune cell populations in tissues. Insights from this approach inform new avenues to selectively target tissue-specific immune populations, functional subsets, and the interactions driving immune cell function within a given tissue^29,33,34^.

## Supplementary Information: Tables

Supplementary Table 1. Xenium custom mouse 350 gene panel

Supplementary Table 2. Gene expression correlation by subtype and spatial axes

Supplementary Table 3. Merscope custom mouse gene panel

Supplementary Table 4. Xenium custom human gene panel

Supplementary Table 5. KS statistics of cell distances.

## Acknowledgments

We would like to thank Pallav Kosuri, Noorsher Ahmed, Clarence Mah, Carlos Marinas, Roy Maimon, Elsa Molina, the Gene Yeo lab, and the ImmGen consortia for helpful discussions and feedback on data analysis.

## Author contributions

Conceptualization: M.R-C., M.H., and A.W.G.

Methodology: M.R-C., A.M., A.F., V.L., B.B., J.C., M.H., and A.W.G.

Investigation: M.R.C., A.F., V.L., K.P.C., G.G., N.E.S., K.K.T., Y.H.L., W.H.W., C.S.I., M.H.

Visualization: M.R-C., M.H., and A.M.

Funding acquisition: G.W.Y., J.C., and A.W.G.

Project administration: M.R-C., M.H., and A.W.G.

Supervision: M.R-C., M.H., and A.W.G.

Writing – original draft: M.R-C., A.M., M.H., and A.W.G.

Writing – review & editing: M.R-C., A.M., M.H., and A.W.G.

## Data and materials availability

The code and data to reproduce the analysis presented in this manuscript are available upon reasonable request.

---

**Extended Data 1, related to Figure 1.**
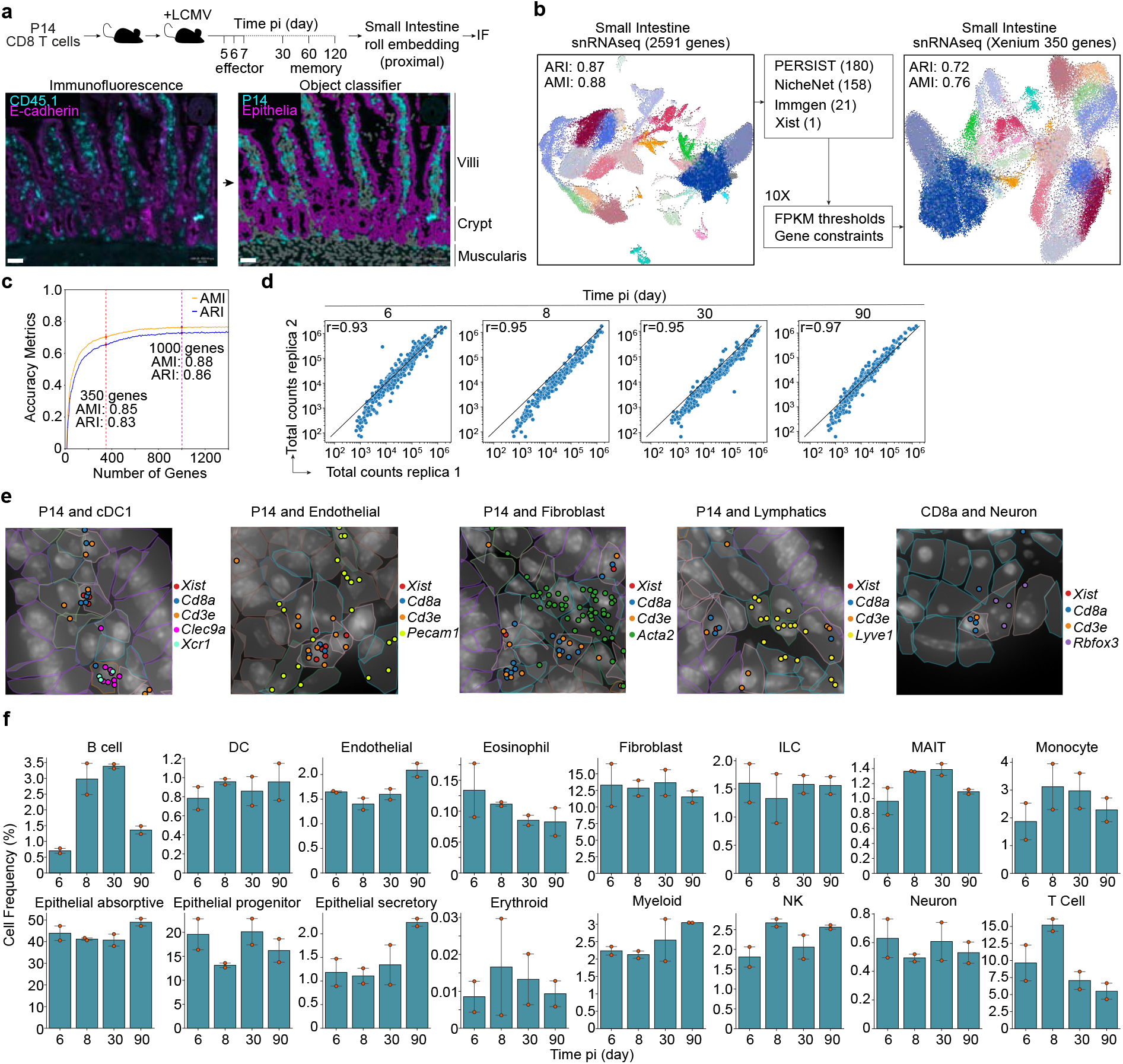
**a**, Schematic of the experimental workflow for mouse takedown at progressing timepoints post LCMV infection (pi) before Immunofluorescence staining. An object classifier in QuPath is used to identify P14 and epithelial cells from IF staining. **b**, Diagram of the methodology used to design the Xenium mouse SI probe panel. Using snRNA-seq data from the mouse small intestine, a 350 gene set was designed to maximize Adjusted Rand Index (ARI) and Adjusted Mutual Information (AMI) scores of classifier-derived Leiden cluster predictions. **c**, Genes least informative for predicting cell type are continuously pruned using recursive feature elimination with ARI and AMI of classifier-derived Leiden cluster predictions calculated at each pruning step. **d**, Pearson residual correlations of total gene abundances between timepoint biological replicates. **e**, Snapshots from a Xenium spatial transcriptomics day 6 small intestine show unique cell types in close spatial proximity. Canonical cell type marker gene transcript positions are colored to show a (from left to right) P14 T Cell (*Xist*^+^*Cd8α*^+^*Cd3e*^+^) and cDC1 (*Clec9a*^+^*Xcr1*^+^), P14 T Cell and Endothelial (*Pecam1*^+^), P14 T Cell and Fibroblast (*Acta2*^+^), P14 T Cell and Lymphatic (*Lyve1*^+^), and Cd8α T Cell (*Cd8α*^+^*Cd3e*^+^) and Neuron (*Rbfox3*^+^). Predicted cell segmentation boundaries are colored by the “Type” annotation. **f**, Cell type frequency percentages across all timepoints. Error bars span the percentages for the two biological replicates per time point.

**Extended Data 2, related to Figure 2.**
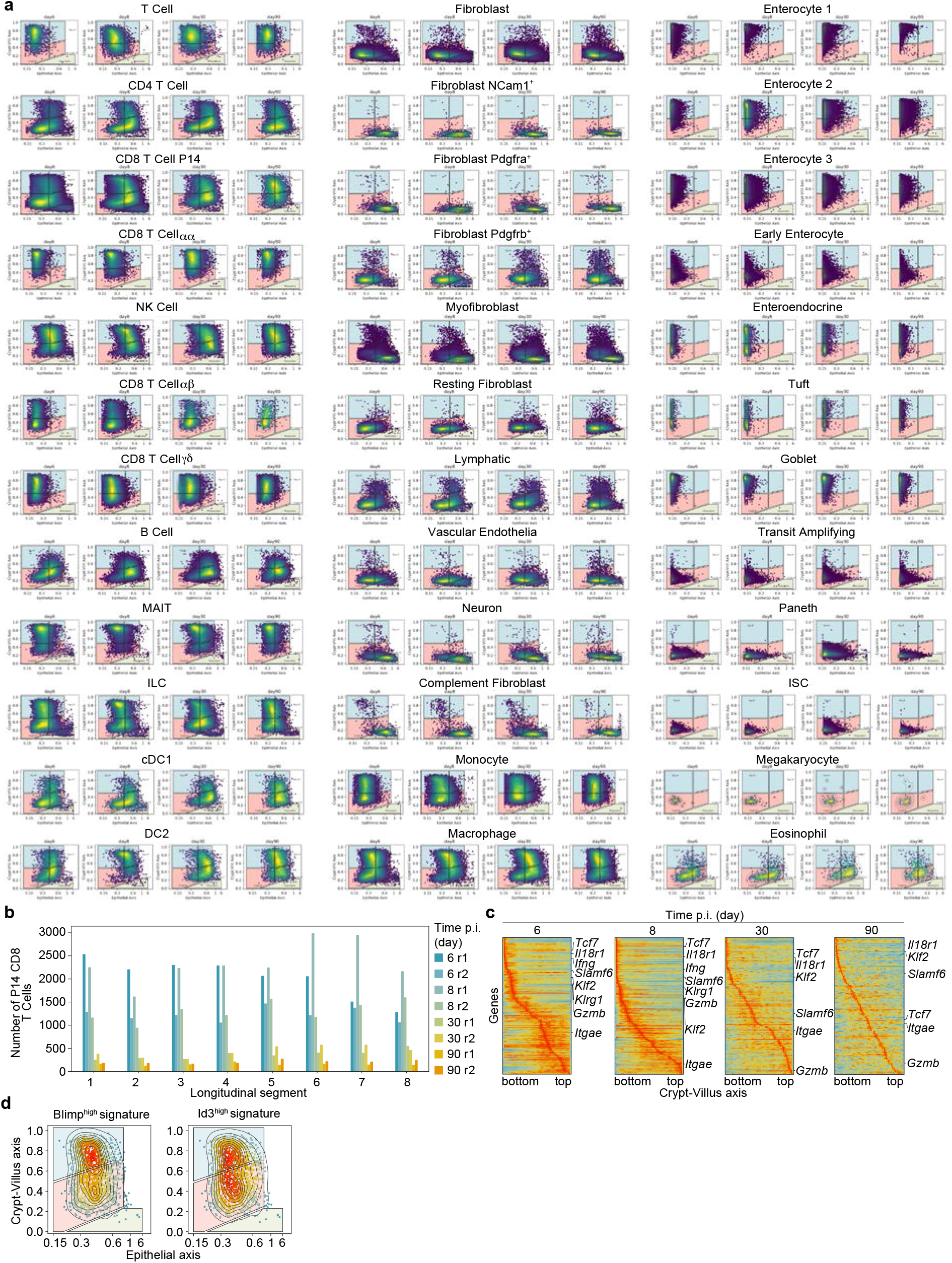
**a**, IMAPs of all cell subtypes from each time point (two biological replicates for each time point), with colored gates dividing the top IE, top LP, crypt IE, crypt LP, and muscularis. **b**, Number of P14 CD8 T cells present in binned segments of equal length of the longitudinal axis for each replicate and time point. **c**, Convolved gene expression of P14 cells along the crypt-villus axis at every timepoint (n = 2 pooled biological replicates). **d**, IMAPs of day 90 P14 CD8 T cells (one of two biological replicates) colored by Blimp1^high^ effector-like and Id3^high^ memory precursor-like signature enrichment derived from Milner et al.

**Extended Data 3, related to Figure 3.**
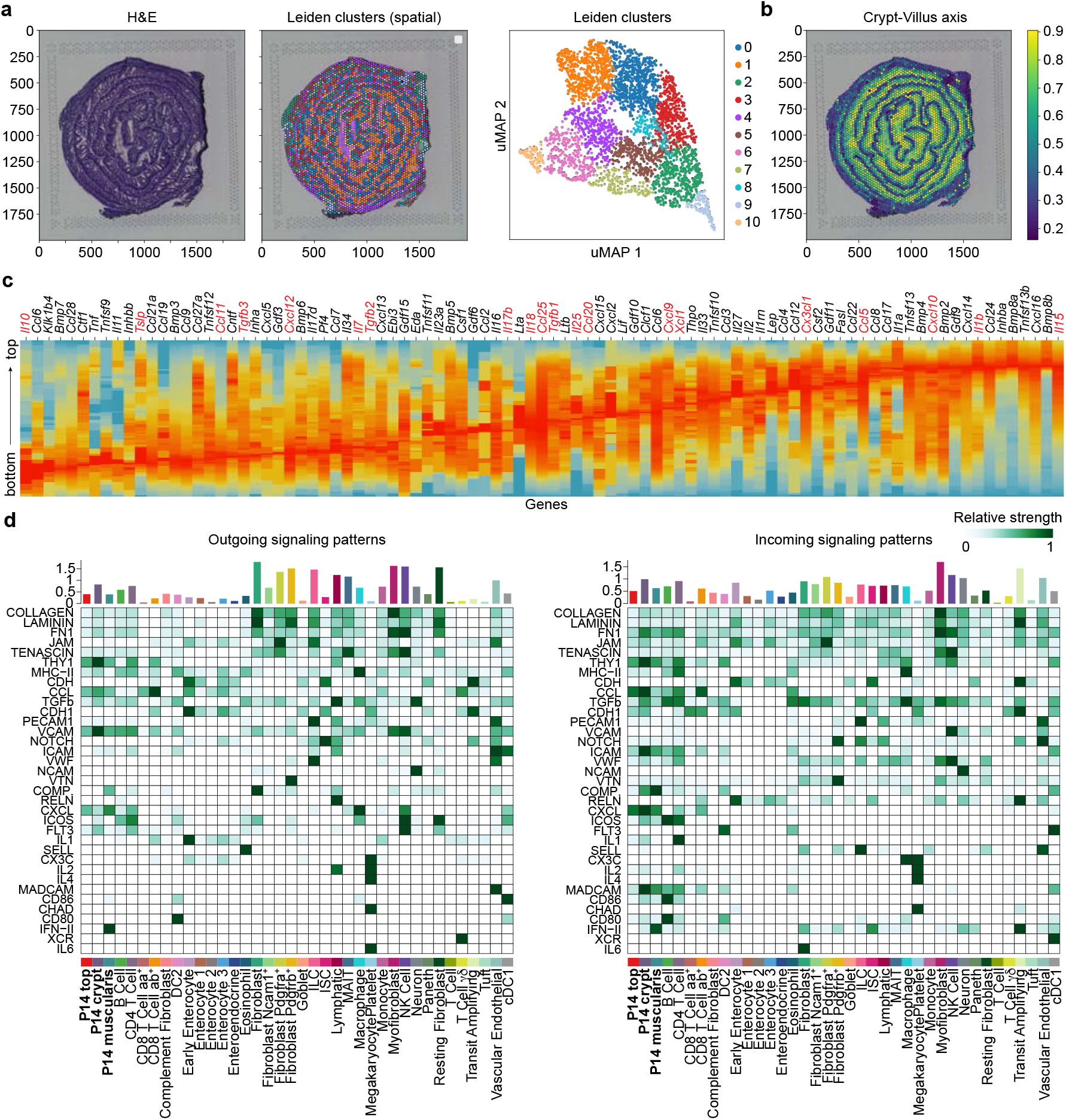
**a**, Overview of Visium-based spatial transcriptomics on day 21 mouse SI tissue - (left) H&E staining and (right) spots Leiden-clustered according to RNA expression and positioned on the tissue. **b**, Visium spots colored by imputed crypt-villus axis values using continuous label transfer after jointly embedding with the mouse Xenium day 30 SI. **c**, Convolved gene expression of cytokines along the imputed crypt-villus axis in the Visium sample. Red labels indicate genes that were included in the Xenium gene panel. **d**, Full version of **Fig 3g**. Strength of outgoing (left), and incoming (right) signals in each signaling pathway across all cell subtypes. P14 CD8 T cells are separated into distinct regional types based on IMAP gating.

**Extended Data 4, related to Figure 4.**
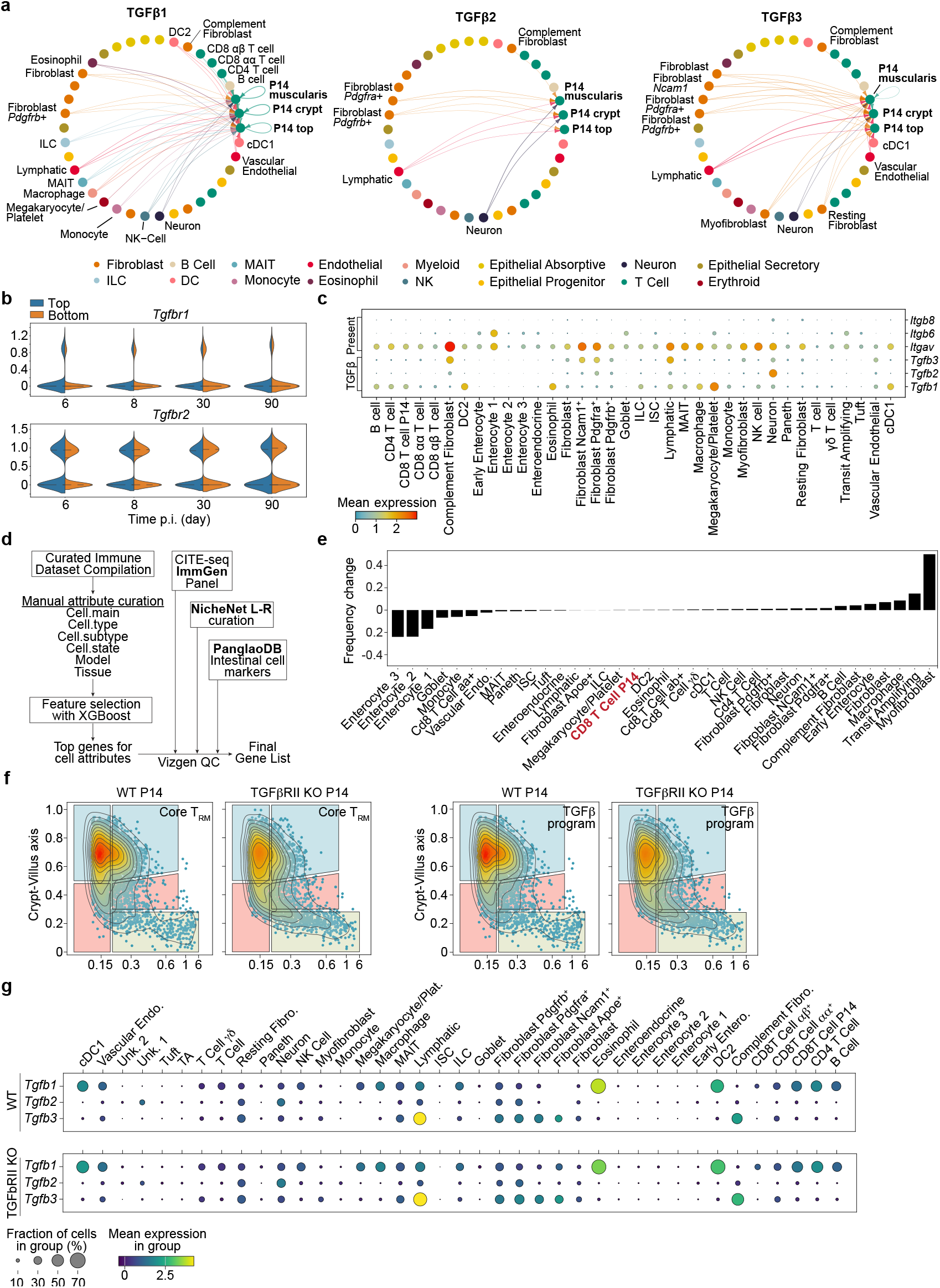
**a**, CellChat circle plots showing top enriched interactions between TGFβ isoform senders and regionally gated P14 receivers across all 8 SI samples (4 timepoints, 2 replicates each). Each cell subtype is represented by a node, and a directed edge is displayed from a top sender subtype to a receiver P14 regional subtype for significant TGFβ sender-P14 interactions. **b**, Violin plots depicting the log-normalized TGFβRI and TGFβRII expression counts within P14s across each timepoint. Violins are plotted with Scanpy, scaled by width, and black dotted lines mark expression quartiles. **c**, The expression of TGFβ isoforms and genes involved in TGFβ presentation from **Fig 4g** as measured by Xenium. **d**, Schematic for MERSCOPE gene panel design process. Most important genes for defining cell types were identified using XGBoost on a group of immune datasets, before adding biologically important genes and filtering out MERSCOPE-incompatible genes. **e**, Change in frequency of each cell type between WT and TGFβR2 KO conditions. Frequency values reflect proportional increases or decreases of TGFβR2 KO cell type counts relative to WT. **f**, IMAPs of WT and TGFβR2 KO P14 T cells colored by enrichment of the core T_RM_ signature from Milner et al, and the TGFβ program derived from Nath et al. **g**, A comparison of TGFβ isoform expression between cell subtypes in WT and TGFβR2 KO.

**Extended Data 5, related to Figure 5.**
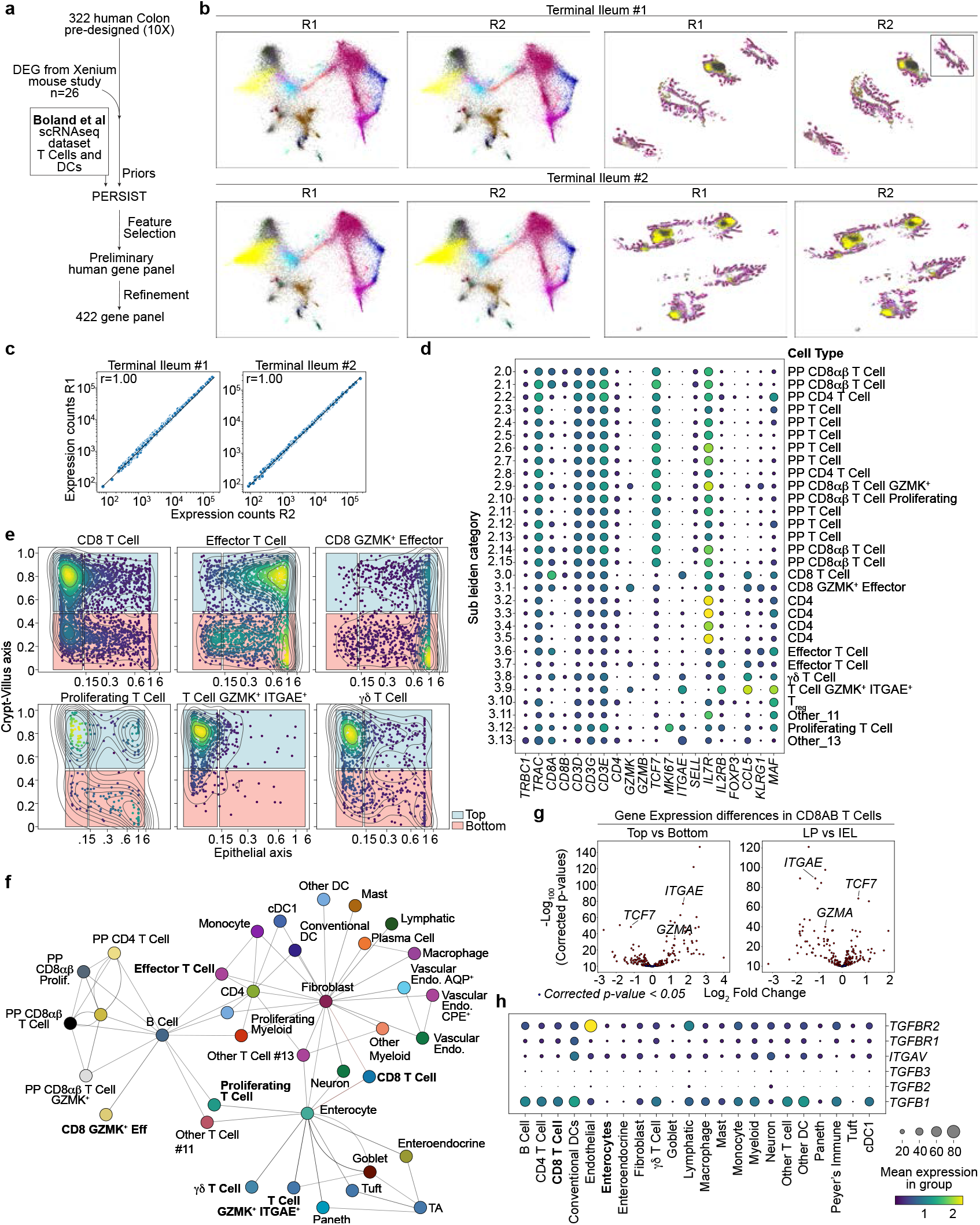
**a**, Schematic for designing the Xenium human SI gene panel. The Xenium base human colon panel was expanded with canonical immune genes, the human homologs of top spatially differentially expressed genes from the Xenium mouse data, and computationally derived genes that best capture the heterogeneity within immune cell types found in scRNA-seq data from Boland et al. **b**, Xenium processed terminal ileum samples divided into two rows corresponding to the two human donors. Adjacent tissue sections were taken from both donors and are positioned side-by-side within the joint MDE embedding (left) and spatially (right). Cells are colored by their annotations in **Fig 5a. c**, Scattered raw gene expression abundances between the technical replicates of both human ileums overlayed with a line of best fit. The Pearson residual correlation coefficient (r) is calculated between the gene abundances of both samples. **d**, Expression of genes used to annotate immune subtypes. Colors of dots indicate the mean expression of the gene in each subcluster, and size of the dots correspond to the percentage of cells in each subcluster expressing the gene. The final cell subtype annotations of each subcluster are shown as y-ticks along the right side of the plot. **e**, IMAP positioning of select T-Cell subtypes within all (n=4) human sections (Peyer’s Patches excluded). Cells are colored by kernel density estimates of their coordinate location within the IMAP. IMAP gates are positioned as in **Fig 5d. f**, Aggregated physical interaction network where edges between nodes represent a normalized Squidpy interaction score lying above a 0.1 threshold (10% of the connections). Nodes are positioned using a Kamada-Kawai layout algorithm on the averaged interaction matrix of all human sections. **g**, Differential expression testing of all genes expressed in at least 5% of human CD8αβ T Cells using diffxpy. A two-tailed Wald test yielded a fold change and adjusted p-value (padj) for each gene (X) between human CD8αβ T cells gated in the crypt versus those gated in the top of the villus, and (X) human CD8αβ T cells gated intraepithelial versus those gated in the lamina propria. All genes are plotted by their log2 fold change and -log100(padj), and significantly differentially expressed genes (padj < 0.05) are colored red. **h**, Expression of TGFβ isoforms and genes involved in TGFβ presentation across cell types after pooling the cells from all human sections (n=4).

## Methods

### Mice

Mice were maintained in specific-pathogen-free conditions at a temperature between 18 °C and 23 °C with 40–60% humidity and a 12h-light and 12h-dark light cycle in accordance with the Institutional Animal Care and Use Committees (IACUC) of the University of California San Diego (UCSD). All mice were of C57BL/6J background and bred at the University of California San Diego (UCSD) or purchased from the Jackson Laboratory. R26Cre-ERT2 (stock no. 008463, Jackson Laboratory), Tgfbr2^fl/fl^ (stock no. 012603, Jackson Laboratory), P14, and CD45.1 congenic mice were bred in-house. Male recipient mice were used for adoptive transfer experiments, and females were used as P14 CD8 T cell donors. To delete floxed alleles using Cre-ERT2, 1 mg of tamoxifen (Cayman Chemical Company) emulsified in 100 μl of sunflower seed oil (Sigma-Aldrich) was administered via intraperitoneal injection (i.p.) for five consecutive days to P14;Cre-ERT2;Tgfbr2^wt^ (WT) and P14; Cre-ERT2;Tgfbr2^fl/fl^ (TGFβRII KO) mice prior to P14 CD8 T cell isolation. All mice were between 1.5 and 6 months old at the time of infection and randomly assigned to experimental groups. No statistical methods were used to pre-determine sample sizes, but our sample sizes are like those reported in previous publications from our laboratory and others. No blinding was performed during mouse experiments. Investigators were not blinded to group allocation during data collection and/or analysis. Mice were fed ad libitum for the specified amount of time. All animal studies were approved by the Institutional Animal Care and Use Committees of UCSD and performed in accordance with UC guidelines.

### Adoptive cell transfer of Naive P14 CD8 T cells and LCMV infection in mice

5×10^4^ female naive P14 CD8 T cells isolated by negative enrichment using MACS magnetic columns and resuspended in PBS were transferred intravenously into congenically distinct male recipient mice. Recipient mice were subsequently infected intraperitoneally with 2×10^5^ plaque-forming units (PFU) of the Armstrong strain of LCMV.

### Preparation of single-cell suspensions for flow cytometry

Isolation of CD8 T cells was performed similarly as described^1^. Small intestine (SI) intra-epithelial lymphocytes (IEL) and lamina propria lymphocytes (LPL) were prepared by removing Peyer’s patches and the luminal contents from the entire SI. The SI was then cut longitudinally and into 1 cm pieces, then incubated at 37°C for 30 minutes in HBSS with 2.1 mg/mL sodium bicarbonate, 2.4 mg/mL HEPES, 8% bovine growth serum, and 0.154 mg/mL of dithioerythritol (EMD Millipore). Collection of the supernatant through a 70 μM constituted the IEL compartment of the SI. The remaining tissue fragments of the SI were further incubated in RPMI with 1.2 mg/mL HEPES, 292 μ/mL L-glutamine, 1 mM MgCl2, 1 mM CaCl2, 5% fetal bovine serum, and 100 U/mL collagenase (Worthington) at 37°C for 30 min. After enzymatic incubation, tissues were filtered through a 70-μm nylon cell strainer (Falcon). Tissue preparations were separated on a 44%/67% Percoll density gradient.

### Sample preparation for histology of the mouse small intestine

<u>For Fresh frozen (FrF)</u> samples, mouse SI were harvested, retaining the proximal-distal orientation. After discarding the first 3 cm proximal section, approximately 10 cm of mouse proximal small intestine was rinsed in ice-cold PBS and the lumen contents flushed with 20 mL of ice-cold PBS using a gavage syringe. The SI was then loaded onto a 3.25mm diameter knitting needle pre-moistened with cold PBS and placed directly on thick blotting paper. MIST^2^ was used as a guide for the scalpel to cut the intestine longitudinally along the knitting needle. The MIST and needle were removed, and the SI was spread open and rolled using a wood autoclaved round toothpick and snap-frozen in OCT in plastic molds for cryosection (Tissue-Tek Cryomold) and kept at -80°C until processing. For <u>Fixed frozen (FiF)</u> samples, the opened cleaned SI were fastened to blotting paper by minute pins in each corner and fixed in 4% paraformaldehyde solution in PBS 4ºC for 16 hours, followed by incubation in 70% ethanol at 4ºC for a minimum of 3 hours. SI samples were then rolled using a wood autoclaved round toothpick and snap-frozen in OCT in plastic molds for cryosection (Tissue-Tek Cryomold) and kept at -80ºC until processing. For <u>Formalin-Fixed Paraffin-Embedded (FFPE)</u> samples, fixed SI were rolled, mounted on 2% agar round molds, and placed on histology cassettes for paraffin embedding.

### Histology and Immunofluorescence staining of fresh frozen mouse tissues

After OCT block equilibration at -20ºC, 10 μm slices were obtained using a cryostat, mounted on glass slides, dried for 20 minutes at - 20ºC, and fixed in ice-cold acetone at -20ºC for 20 minutes. After fixation, slides were dried briefly at room temperature and stored at -80ºC until stained or used immediately. For staining, slides were equilibrated at room temperature, washed in 4ºC PBS twice for five minutes, blocked in serum-free blocking reagent (Dako) ON at 4°C, followed by staining with CD45.1-AF594 (BioLegend, clone A20, Reference #110756) and E-cadherin-APC (BioLegend, clone DECMA-1, Reference #147312), and CD8a-FITC (BioLegend, clone 53-6.7, Reference #35-0081-U500) diluted in Antibody diluent solution (Dako, S080983-2) overnight at 4ºC, stained with DAPI, and mounted with coverslips using Vectashield Vibrance Antifade mounting media (VectorLabs, H-1700). Images were acquired on an Olympus VS200 Slide Scanner (UCSD Microscopy CORE) or on a ZEISS LSM700 confocal microscope. Quantifying P14 CD8 T cell distances for IMAP representation over time was done using a groovy script on QuPath.

### Single Nuclear RNA Sequencing (snRNA-seq) of mouse SI

Female CD45.1^+^ CD8^+^ P14 T cells were adoptively transferred into male CD45.2^+^ recipients (1×10^5^ cells/mouse) 30 minutes prior to infection with LCMV-Armstrong. At 28 days post infection, mice were euthanized and small intestine was dissected, flayed, and washed in cold PBS; Peyer’s patches were excised from the small intestine. The small intestine was divided into three equal sections designated as the proximal, middle, and distal small intestine. Tissue sections were cut into ∼3 mm pieces and flash-frozen in liquid nitrogen for 2 min. Nuclei isolation was performed with 10x Genomics Chromium Nuclei Isolation Kit per the manufacturer’s instructions. Briefly, 30-50 mg of flash-frozen tissue per sample was dissociated with a pestle, incubated for 10 minutes on ice, and washed. Dissociated tissue was passed through a Nuclei Isolation Column, and flowthrough nuclei were washed in Debris Removal Buffer and Wash and Resuspension Buffer. Nuclei were quantified with Nexcelom Bioscience Cellometer. For maximum targeted recovery, 40,000 nuclei per sample were loaded for GEM generation. Samples were processed via the Chromium Next Gem Single Cell 3’ HT Dual Index v3.1 protocol and sequenced to a depth of 550 million read pairs per sample (∼23,000 read pairs per nucleus) on a NovaSeq 6000 system (Illumina).

### Spatial transcriptomics analysis using whole-genome spatial transcriptomics (Visium, 10X)

CD45.1^+^ CD8^+^ P14 T cells were adoptively transferred into CD45.2^+^ recipients (1×10^5^ cells/mouse) 30 minutes prior to infection with LCMV-Armstrong. At 21 days post infection, mice were euthanized, and their small intestines divided into proximal and distal sections. The small intestine sections were rolled up from distal-to-proximal into “swiss rolls.” The swiss rolls were embedded in Optimal Cutting Temperature (OCT) (Sakura Finetek) and flash frozen in 2-methyl butane (Sigma Aldrich) over dry ice. Tissue blocks with an RNA Integrity Number (RIN) 7 were cryosectioned at 20μm thickness and mounted onto the Visium Spatial slide (10x Genomics). The slide was fixed in methanol (Millipore Sigma) then Hematoxylin & Eosin (H&E) stained (Agilent) according to the Visium Spatial protocol (CG000160, 10x Genomics). Brightfield images were acquired using the BZX-700 Fluorescent Microscope (Keyence) at 20X magnification. Libraries were constructed according to the demonstrated Visium Spatial Gene Expression protocol (CG000239, 10x Genomics) with an optimized permeabilization time of 24 minutes. Pooled libraries were sequenced on a NovaSeq 6000 system (Illumina).

### Spatial transcriptomics analysis using Multiple Error-Robust Fluorescence in situ Hybridization (MERFISH)

Fresh frozen tissue was sectioned according to standard histology procedures to a thickness of 10 μm. The sections were adhered to the MERSCOPE slides (Vizgen, #20400001) coated with fluorescent beads by storing them in the cryostat at -20° C for at least 5 minutes. The samples were fixed in 5 mL of fixation buffer containing 4% paraformaldehyde in 1x Phosphate Buffered Saline (PBS) that was preheated to 47° C and incubated for 30 min at 47° C, according to the MERSCOPE Quick Guide Modified Fixation for Fresh Frozen Samples. The samples were then washed 3 times with 5mL PBS, 5 minutes each time. The samples were permeabilized in 5 mL 70% ethanol at 4° C in parafilm-sealed dishes overnight and stored in these same conditions for up to a month. Samples were then prepared according to Vizgen’s protocols, starting from the cell boundary protein staining step. The samples were hybridized with a custom 500-gene panel that included 5 sequential genes, as well as several blank barcodes that do not encode a gene and used for background signal measurement. To clear the samples of lipids and proteins that interfere with imaging, 5 mL of Clearing Premix (Vizgen #20300003) was mixed with 100 μL of Proteinase K for each sample, and the samples were placed at 47° C in a humidified incubator overnight (or for a maximum of 24 hours), and then moved to 37° C. The samples were stored in the Clearing solution in the 37° C incubator prior to imaging for up to a week. The samples were imaged on the MERSCOPE according to the MERSCOPE Instrument User Guide. Seven 1.5 μm-thick z planes were imaged for each field of view at 60x magnification. Images were decoded to RNA spots with xyz and gene id using Vizgen’s Merlin software. Cell segmentation was performed using the Cellboundary algorithm, relying on the Cellboundary 2 stain and DAPI nuclear seeds.

### Spatial transcriptomics analysis using 10X Xenium

FFPE tissues were sectioned to a thickness of 5 μm onto a Xenium slide, followed by deparaffinization and permeabilization following the 10X user guides CG000578 and CG000580. Probe hybridization, ligation, and amplification were done following the 10X user guide CG000582. In short, probe hybridization occurred at 50 °C overnight with a probe concentration of 10 nM using a custom gene panel designed to detect 350 different mRNAs. After stringency washing to remove un-hybridized probes, probes were ligated at 37 °C for two hours. During this step, a rolling circle amplification (RCA) primer was also annealed. The circularized probes were then enzymatically amplified (two hours at 37 °C), generating multiple copies of the gene-specific barcode for each RNA binding event. After washing, background fluorescence was quenched chemically. Sections were placed into an imaging cassette to be loaded onto the Xenium Analyzer instrument following the 10X user guide CG000584.

### Spatial Data Processing

For 10x Xenium spatial transcriptomics data, nuclei were segmented using a fine-tuned Cellpose^5^ model on max-projected DAPI staining images. Baysor^6^ was used to predict cell boundary segmentations using the Cellpose nuclei segmentations as a prior segmentation for 10x Xenium or the cellboundary 2 segmentation for MERSCOPE respectively and transcripts as input, with parameters prior-segmentation-confidence = 0.95 and 0.9 respectively, and min-molecules-per-cell set to the median nucleus transcript count (see https://github.com/Goldrathlab/Spatial-TRM-paper#preprocessing). Baysor segmentations containing no nuclei were filtered out, and segmentations containing multiple nuclei were split by assigning transcripts to the nearest nucleus centroid within the segmentation boundary. All cells with n < 8 nuclear transcripts, n < 20 total transcripts, or n > 800 total transcripts were filtered out before downstream processing. To integrate spatial replicates into a joint embedding, scVI was used with n_layers = 2, and n_latent = 30. The joint embedding was project into 2D space using scVI.model.utils. mde. Leiden clustering was performed on the scVI learned embeddings using scanpy.tl.leiden with resolution=1, and every Leiden cluster was further subclustered at resolution = 1.2. Celltypist^7^ and GeneFormer^31^ were used for a first-pass cell type assignment, with further manual refinement based on the expression of cell type marker genes to define cell types in a Class > Type > Subtype hierarchy. Anndata^4^ format was used for all further processing. To align histology images with Xenium spatial coordinates, we utilized an OpenCV ORB object to detect key points in the DAPI channel of both histology and Xenium images. These key points were then matched using an OpenCV DescriptorMatcher, enabling the computation of a homography matrix based on the top matches using cv2.findHomography. Subsequently, histology images across all channels underwent warping using this homography matrix with cv2.warpPerspective. To align H&E images with Xenium spatial coordinates, we trained a pix2pix generative adversarial network^8^ to predict DAPI images from H&E as an intermediate state before finding key points and matching, as previously mentioned. To visualize mouse transcriptional signatures onto human datasets, all (n = 8) mice time course samples were used to find the top 15 differentially expressed genes (Scanpy rank_ genes_groups and method = ‘wilcoxon’) between P14s gated to the crypt and P14s gated to the top of villi. Human homologs of these 15 genes are defined as UCell signatures and mapped to human CD8 AB+ T Cells. Human CD8 AB+ T Cells are positioned on IMAPs and colored by their enrichment of the (left) top mouse signature and (right) crypt mouse signature. All code to analyze the spatial datasets can be found on https://github.com/Goldrathlab/Spatial-TRM-paper.

### Histological staining of mouse intestinal tissue after Xenium analysis

Post Xenium run, slides were kept hydrated in PBS-T (0.05% tween-20 in PBS) at 4ºC. For post-Xenium immunofluorescent staining, PBS-T was removed and samples were blocked using universal blocking reagent CAS-Block, and stained with anti-CD8α (Abcam) overnight at 4ºC, followed by 3 washes of PBS. Anti-rabbit AF594 secondary antibody in Dako antibody diluent was then added for 1 hour at room temperature in the dark, followed by 3 washes of PBS. The slides were then stained by WGA-FITC followed by DAPI staining, and then mounted with a coverslip using Vectashield mounting medium. Slides are dried for 1 hour at room temperature in the dark prior to imaging with Olympus VS200 Slide Scanner at 20X. Slides were then soaked in PBS at 4ºC overnight to dismount coverslip and subsequently washed 3 times in PBS and twice in ddH_2_O before proceeding with H&E staining. The coverslip was mounted using xylene-based mounting media (Cytoseal XYL Epredia), slides were dried for 1 hour, and imaged used VS200 Slide Scanner at 20x.

### Human samples

Deidentified human FFPE samples from healthy subjects were acquired from the San Diego Digestive Diseases Research Center (SDDRC). 5 μm slices were obtained using a microtome, deparaffinized, and H&E stained by common histology practices or processed according to the 10X Xenium protocol.

### Human subjects and ethical statement

The Human Research Protection Programs at the University of California, San Diego reviewed and approved the protocol, including a waiver of consent. Submucosal ileal biopsies were obtained from patients who underwent colonoscopies to rule out inflammatory bowel diseases. Ileal biopsies were evaluated by a pathologist and found to be normal without histologic inflammation. Samples were de-identified and processed for the study.

### Gene panel design for probe-based spatial transcriptomics profiling of mouse small intestine

For the Xenium mouse 350 gene panel, 79 small intestine canonical cell type marker genes were compiled from existing literature, and Xist, a marker for transferred female P14 CD8 T cells. An additional 158 genes from a Nichenet database^9^ of ligand-receptor pairs were included. Next, supervised PERSIST^10^ was run on an immune-enriched gut scRNA-seq dataset^11^ with the previously compiled set of genes as prior information, adding an additional 70 genes. Lastly, supervised PERSIST was run on a SI scRNA-seq dataset to capture 59 cell type marker genes for 350 total targets. To create the Xenium human 422 gene panel, we created a base set of canonical immune marker genes, ligand-receptor pairs, spatially differentially expressed genes within mouse P14 CD8 T cells, and the 10X Genomics base human colon panel totaling 343 genes. Using this set as prior information to PERSIST, and a reference human immune cell scRNA-seq dataset^12^, unsupervised PERSIST filled in the remaining 79 genes. To create the 494 gene panel for MERSCOPE, we compiled 18 published bulk RNAseq datasets profiling different immune populations in different disease settings^13–27^, including the ImmGen RNAseq database (immgen.org), and manually curated metadata attributes for each sample for the following categories: cell.main, cell.type, cell.subtype, cell.state, model, and tissue. The annotated integrated dataset compilation was used as input for Feature Selection with XGBoost^28^. Top genes within each attribute were incorporated as the panel backbone. In addition, gene markers of intestinal cell markers from PanglaoDB and Haber et al 2017^29^, ImmGen CITE-seq protein markers, and 159 genes from the ligand-receptor database NicheNet^9^ were added. 494 final genes passed Vizgen’s quality check filtering for transcript length and expression levels.

### Defining structural axes in spatial transcriptomics datasets of the small intestine

To calculate the <u>longitudinal axis</u>, a multi-line segment was initially labeled across the base of the basal membrane, starting from the outermost section of the roll. For each cell, the nearest neighbor was calculated from the set of all locations on the multiline segment positioned closer to the center of the SI roll. The relative position of the nearest neighbor along the length of the entire multi-line segment was used as the longitudinal position. In datasets with atypical morphology, longitudinal axis values for each cell were predicted using a deep neural network trained on a feature space of transcriptional neighborhood decomposition latent factors and multi-line segments marking the base of the basal membrane, the top of each villus, and the middle of each villus. Transcriptional neighborhood decomposition was performed using non-negative matrix factorization on a matrix of the summed transcript count values for the 10 nearest neighbors of each cell, to create a transformed data matrix W with 15 latent factors. To calculate the <u>crypt-villus axis</u>, a fine-tuned Cellpose model was used to segment villi based on WGA staining in Xenium samples and cell spatial positions in MERSCOPE samples. The distance between each cell and the base of the basal membrane was calculated as before, and these values were standard scaled among cells in the same segmented villus. For datasets with poorly defined morphology, crypt-villus axis values for each cell were predicted using a deep neural network trained on a feature space of epithelial and stromal transcriptional neighborhood decomposition latent factors. Transcriptional neighborhood decomposition was performed by fitting a non-negative matrix factorization model on a matrix of the summed transcript count values for the (10 – mouse, 30 – human) the nearest epithelial and stromal neighbors of each cell in all datasets to avoid the influence of variability in immune populations at different infection states. Crypt-villus axis predictions for each cell were smoothed over their nearest 150 neighboring cell predictions in human data. To calculate <u>the epithelial axis</u>, the mean distance from each cell to the five nearest epithelial cells was divided by the mean distance to the five nearest cells of any cell type. The resulting values were standard scaled and clipped at an upper bound to align epithelial distances between the villus and basal membrane.

### Generation of Immune Allocation Plots (IMAPs) and transcriptional IMAPs

The epithelial iMAP axis values are computed through a biexponential transformation applied to the clipped epithelial axis values across all cells. Each cell is positioned on the IMAP by its corresponding crypt-villus axis and transformed epithelial axis values. Density within the scattered point cloud is visualized using color mapped scipy.stats.gaussian_kde values, with density lines overlaid using seaborn.kdeplot for enhanced clarity and interpretation. Gate boundaries were manually drawn to distinguish the muscularis, villus crypt, and villus top by observing the IMAP locations of cell types known to localize to each region. Transcriptional IMAPs were colored by adding an array of gene expression counts as a point weight parameter to the scipy.stats. gaussian_kde function. Similarly, gene signature IMAPs were colored using the squared UCell^30^ signature enrichment scores as the point weight parameter. In human IMAPs, signature and gene expression point weights were squared to overcome bias in CD8αβ^+^ physical density in IMAP visualization.

### Statistical analysis

In **Fig. 2e**, significance cutoffs are set at absolute Spearman ρ > 0.05, an arbitrary threshold used to highlight differences between axes. The Spearman coefficients for each gene and their corresponding p-values are documented in **Supplementary Table 2** across (n=87387) P14 T cells. In **Fig. 4d 4d**, the top 4 differentially expressed genes per condition across P14 cells (n=4135 WT, n=4161 TGFβR2 KO) are calculated using wilcoxon testing with Benjamini-Hochberg correction in the function scanpy.tl.rank_genes_group. In **Fig. 4e and 4f**, a nonparametric two-sample Kolmogorov-Smirnov (KS) statistic is used to calculate significance of difference between P14 cell type proximity distributions in WT and TGFβR2 KO conditions. A cutoff of similarity is arbitrarily positioned at a KS statistic of 0.08, corresponding to a corrected p-value of ∼1e12 – 1e10 (KS p-values vary with number of samples in the compared distributions). KS tests are documented in **Supplementary Table 5**. In Extended Figure 5g, human gene expression counts were log-normalized before differentially expressed genes were calculated between human CD8αβ T cells gated to different regions using a two-tailed Wald test from the Python package diffxpy. P-values were adjusted using a Benjamini-Hochberg correction. Error bars in all figures are created using standard error of the mean.

## Data availability

The data to reproduce the analysis presented in this manuscript is available upon reasonable request.

## Code availability

The code to reproduce the analysis presented in this manuscript is available upon reasonable request.

